## Supplementary Materials for "VASCilia (Vision Analysis StereoCilia): A Napari Plugin for Deep Learning-Based 3D Analysis of Cochlear Hair Cell Stereocilia Bundles"

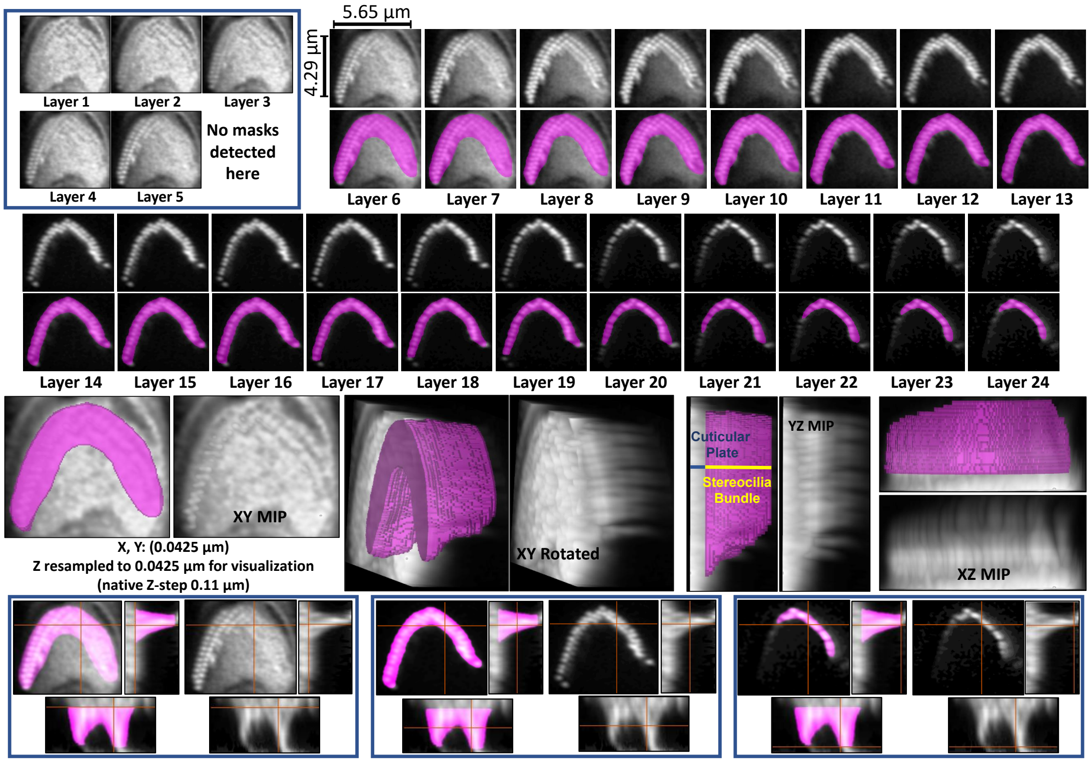


**(a)** Upright bundle rotated into the $z$ plane with strong cuticular-plate signal. Masks follow stereocilia through $z$ and stop where the signal disappears; depth is evident in the 3D and YZ views.

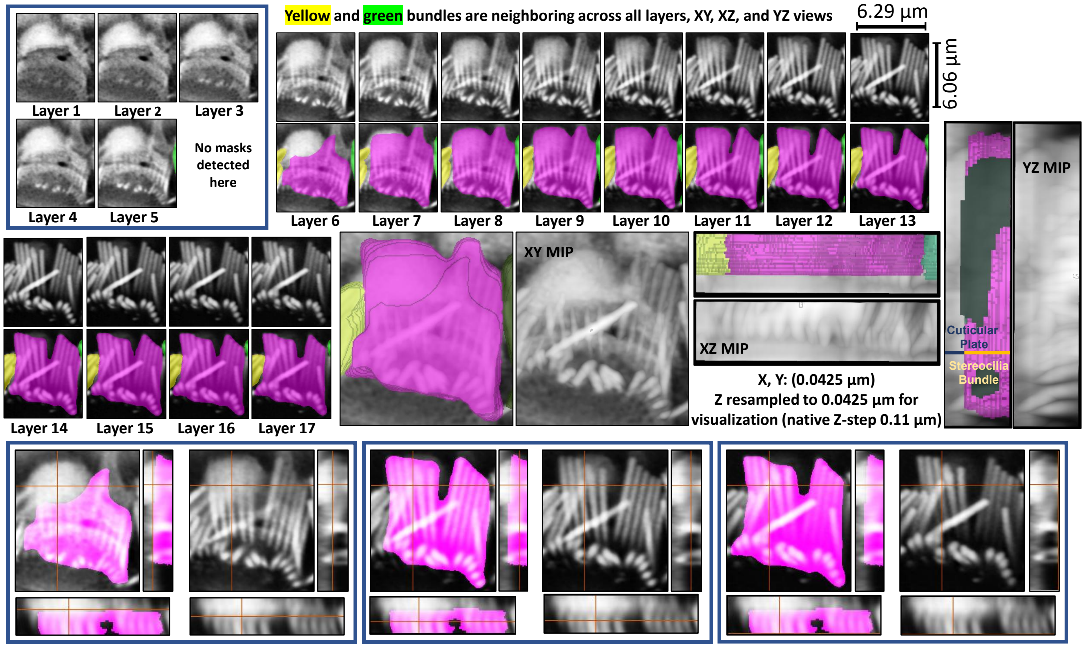


**(b)** Flattened, overlapping bundle. Masks appear where stereocilia are clearly visible (e.g., layer 6) and are absent in previous layers with only background signal, illustrating selective detection.

**Figure 20.** Representative P21 Apex stereocilia segmentations for an upright bundle and a flattened bundle, shown across all $z$ planes with overlays and 3D views. Segmentation depth follows the stereocilia signal and excludes the cuticular plate unless stereocilia signal is clearly present, showing how the method responds to different depths and orientations.

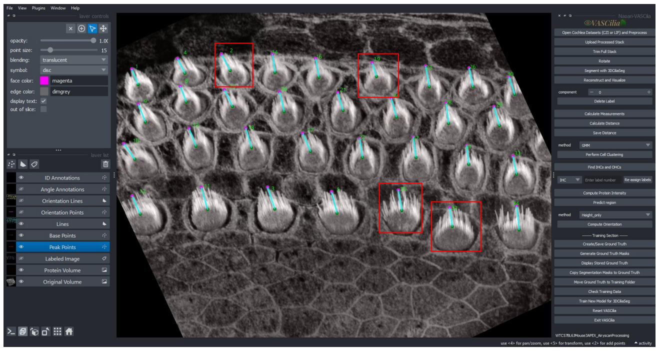

**(a)** Two IHCs and two OHCs from the apical cochlear turn of a P5 WT mouse

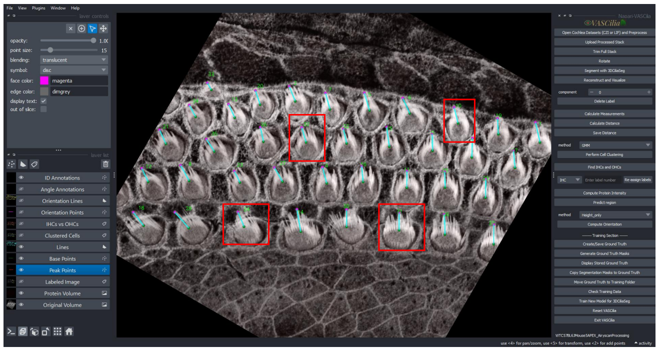

**(b)** Two IHCs and two OHCs from the apical cochlear turn of a P5 WT mouse

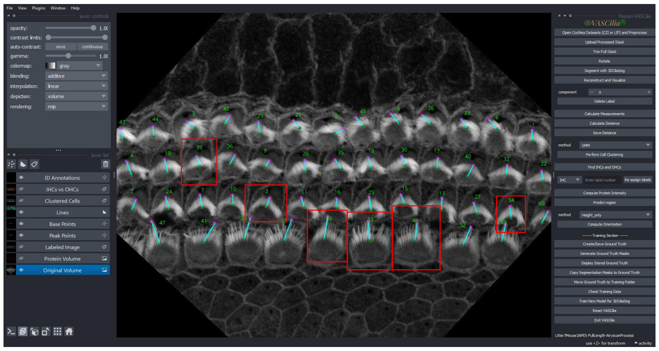

**(c)** Three IHCs and three OHCs were selected from the apical cochlear turn of P5 *Eps8 KO* mouse injected with an AAV–Anc80L65 vector expressing GFP–*Eps8*.

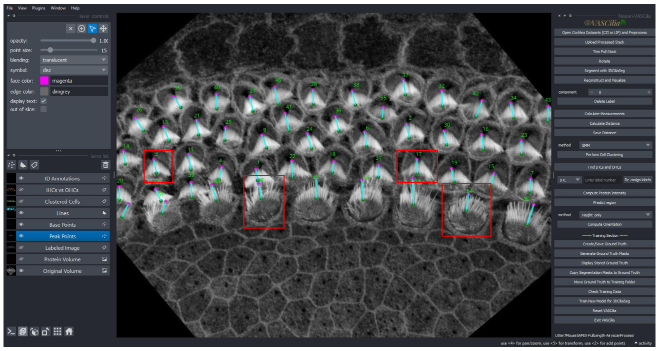

**(d)** Two IHCs and two OHCs were selected from the apical cochlear turn of P5 *Eps8 KO* mouse injected with an AAV–Anc80L65 vector expressing GFP–*Eps8*

**Figure 21.** Full $z$-stacks used to derive the measurement crops summarized in Table 4. Each panel shows the apical cochlear turn; red boxes mark the inner hair cells (IHCs) and outer hair cells (OHCs) from which the crops were taken.

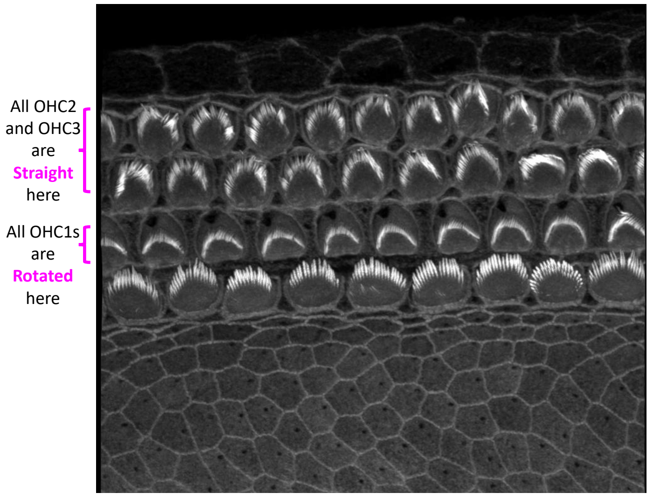


**(a)** Stack from P5 mouse: OHC2 and OHC3 appear straight; OHC1 is rotated.

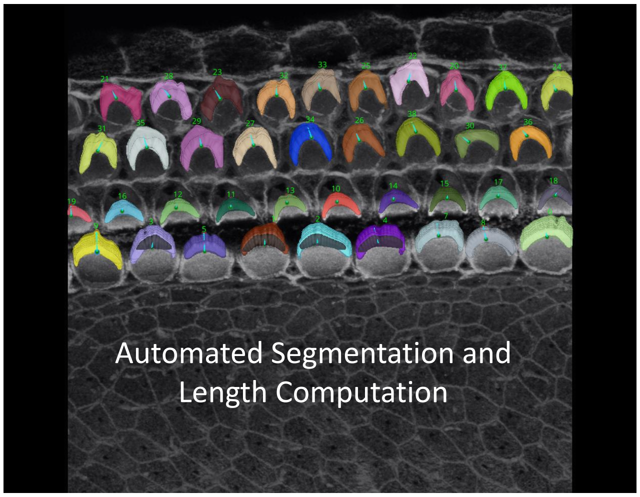


**(b)** Automatic segmentation for the stack.

| ID | Distance | CLass | Depth |
|---|---|---|---|
| 1 | 2.79315 | IHC | 8 |
| 2 | 2.660582 | IHC | 9 |
| 3 | 2.955217 | IHC | 8 |
| 4 | 2.540946 | IHC | 9 |
| 5 | 2.912053 | IHC | 8 |
| 6 | 2.619024 | IHC | 10 |
| 7 | 2.000707 | IHC | 8 |
| 8 | 2.116468 | IHC | 6 |
| 9 | 3.076384 | IHC | 8 |
| 10 | 1.486676 | OHC1 | 10 |
| 11 | 1.586737 | OHC1 | 10 |
| 12 | 1.563191 | OHC1 | 10 |
| 13 | 1.452829 | OHC1 | 10 |
| 14 | 1.550992 | OHC1 | 10 |
| 15 | 1.812385 | OHC1 | 10 |
| 16 | 1.75457 | OHC1 | 10 |
| 17 | 1.65755 | OHC1 | 10 |
| 18 | 1.693337 | OHC1 | 9 |
| 19 | 1.15153 | OHC1 | 5 |
| 20 | 1.939306 | OHC3 | 5 |
| 21 | 1.69437 | OHC3 | 5 |
| 22 | 1.87094 | OHC3 | 5 |
| 23 | 2.089284 | OHC3 | 5 |
| 24 | 1.458561 | OHC3 | 5 |
| 25 | 1.570248 | OHC3 | 5 |
| 26 | 1.678279 | OHC2 | 5 |
| 27 | 1.739153 | OHC2 | 5 |
| 28 | 1.666923 | OHC3 | 5 |
| 29 | 1.653849 | OHC2 | 5 |
| 30 | 1.666923 | OHC2 | 5 |
| 31 | 2.082346 | OHC2 | 5 |
| 32 | 1.554618 | OHC3 | 5 |
| 33 | 1.480717 | OHC3 | 5 |
| 34 | 1.725054 | OHC2 | 5 |
| 35 | 1.881545 | OHC2 | 5 |
| 36 | 1.488029 | OHC2 | 5 |
| 37 | 1.69437 | OHC3 | 5 |
| 38 | 1.716117 | OHC2 | 5 |

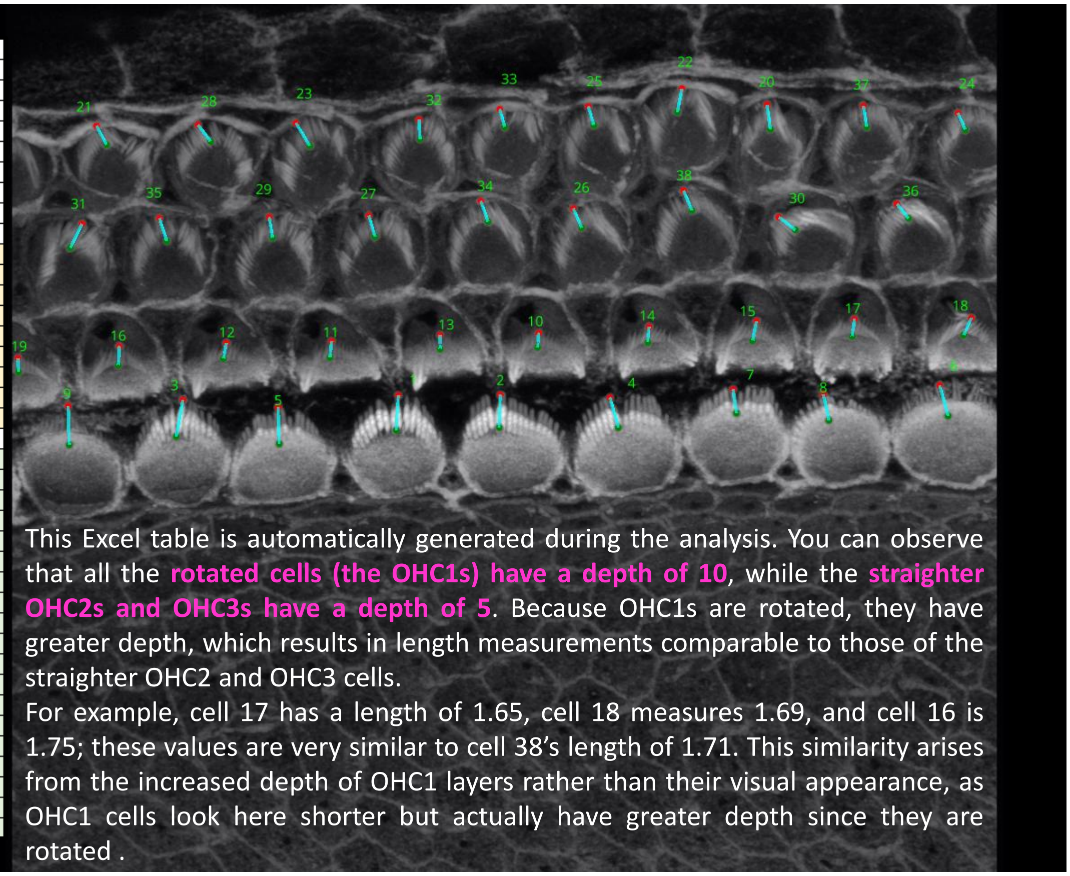


**(c)** Length computation and measurements.

**Figure 22.** Straight vs. rotated (P5): raw stack, automatic segmentation, and length readouts. Because measurements are computed in 3D, the plugin captures bundle depth and robustly handles both straight and rotated bundles—capabilities not achievable with 2D measurements.

| Tonotopic_KO_WT_Class | mean_height | std_height | median_height | count |
|---|---|---|---|---|
| WT_Base_IHC | 3.03 | 0.41 | 2.91 | 26 |
| WT_Middle_IHC | 3.27 | 0.46 | 3.24 | 28 |
| WT_Apex_IHC | 3.90 | 0.55 | 4.01 | 27 |
| KO_Base_IHC | 1.97 | 0.33 | 1.91 | 26 |
| KO_Middle_IHC | 2.07 | 0.24 | 2.05 | 26 |
| KO_Apex_IHC | 2.50 | 0.58 | 2.37 | 26 |
| WT_Base_OHC | 2.29 | 0.27 | 2.32 | 81 |
| WT_Middle_OHC | 2.52 | 0.40 | 2.47 | 85 |
| WT_Apex_OHC | 2.94 | 0.53 | 2.94 | 100 |
| KO_Base_OHC | 1.46 | 0.20 | 1.44 | 77 |
| KO_Middle_OHC | 1.54 | 0.23 | 1.51 | 88 |
| KO_Apex_OHC | 1.91 | 0.38 | 1.90 | 98 |

**Table 7.** Bundle height summary by genotype (WT/KO), tonotopic region (Base, Middle, Apex), and cell type (IHC/OHC), related to Fig. 9a. Values are in $\mu$m; each row reports mean, SD, median, and sample size (N). Row order matches the violin plot. Totals across all groups: IHC $N = 159$, OHC $N = 529$, overall $N = 688$.

| KO_WT | Cell | Pair (level1–level2) | n1/n2 | mean1 | mean2 | Δ (95% CI) | $g$ | $p_{\text{adj}}$ (Holm) | Status |
|---|---|---|---|---|---|---|---|---|---|
| WT | IHC | Base–Middle | 26/28 | 3.03 | 3.27 | −0.24 (−0.48, −0.01) | −0.56 | 0.0425 | Significant |
| WT | IHC | Middle–Apex | 28/27 | 3.27 | 3.90 | −0.63 (−0.90, −0.36) | −1.23 | $5.35 \times 10^{-5}$ | Significant |
| WT | IHC | Base–Apex | 26/27 | 3.03 | 3.90 | −0.87 (−1.14, −0.61) | −1.78 | $8.31 \times 10^{-8}$ | Significant |
| WT | OHC | Base–Middle | 81/85 | 2.29 | 2.52 | −0.23 (−0.34, −0.13) | −0.68 | $1.86 \times 10^{-5}$ | Significant |
| WT | OHC | Middle–Apex | 85/100 | 2.52 | 2.94 | −0.42 (−0.55, −0.28) | −0.87 | $1.42 \times 10^{-8}$ | Significant |
| WT | OHC | Base–Apex | 81/100 | 2.29 | 2.94 | −0.65 (−0.77, −0.53) | −1.50 | $8.70 \times 10^{-20}$ | Significant |
| KO | IHC | Base–Middle | 26/26 | 1.97 | 2.07 | −0.10 (−0.26, 0.06) | −0.35 | 0.209 | Not significant |
| KO | IHC | Middle–Apex | 26/26 | 2.07 | 2.50 | −0.43 (−0.68, −0.18) | −0.95 | 0.00271 | Significant |
| KO | IHC | Base–Apex | 26/26 | 1.97 | 2.50 | −0.54 (−0.80, −0.27) | −1.11 | $6.33 \times 10^{-4}$ | Significant |
| KO | OHC | Base–Middle | 77/88 | 1.46 | 1.54 | −0.08 (−0.15, −0.01) | −0.37 | 0.0164 | Significant |
| KO | OHC | Middle–Apex | 88/98 | 1.54 | 1.91 | −0.36 (−0.45, −0.28) | −1.15 | $3.69 \times 10^{-13}$ | Significant |
| KO | OHC | Base–Apex | 77/98 | 1.46 | 1.91 | −0.45 (−0.53, −0.36) | −1.41 | $8.55 \times 10^{-18}$ | Significant |

**Table 8.** Pairwise tonotopic contrasts in bundle height by genotype (WT/KO) and cell type (IHC/OHC), related to Fig. 9a. Within each stratum we compare Base–Middle, Middle–Apex, and Base–Apex using Welch two-sided $t$-tests. Reported are sample sizes ($n_1/n_2$), group means (mean1/mean2, in $\mu$m), the difference $\Delta = \text{mean}_1 - \text{mean}_2$ with 95% CI, Hedges' $g$, and Holm-adjusted $p$-values. Negative $\Delta$ and $g$ indicate level2 > level1. All contrasts are significant at $p_{\text{adj}} < 0.05$ except KO IHC Base–Middle.

| Tonotopic_KO_WT_Class | mean_value | std_value | median_value | count |
|---|---|---|---|---|
| WT_Base_IHC | 0.70 | 0.27 | 0.84 | 26 |
| WT_Middle_IHC | 0.24 | 0.13 | 0.18 | 28 |
| WT_Apex_IHC | 0.17 | 0.11 | 0.15 | 27 |
| KO_Base_IHC | 0.51 | 0.14 | 0.55 | 26 |
| KO_Middle_IHC | 0.33 | 0.09 | 0.32 | 26 |
| KO_Apex_IHC | 0.29 | 0.24 | 0.25 | 26 |
| WT_Base_OHC | 0.42 | 0.12 | 0.44 | 81 |
| WT_Middle_OHC | 0.16 | 0.09 | 0.13 | 85 |
| WT_Apex_OHC | 0.12 | 0.08 | 0.09 | 100 |
| KO_Base_OHC | 0.28 | 0.11 | 0.24 | 77 |
| KO_Middle_OHC | 0.26 | 0.09 | 0.27 | 88 |
| KO_Apex_OHC | 0.14 | 0.11 | 0.11 | 98 |

**Table 9.** Normalized fluorescence intensity (unitless) by genotype (WT/KO), tonotopic region (Base, Middle, Apex), and cell type (IHC/OHC), ordered to match the violin plot (Fig. 9b). Each row reports the mean, SD, median, and sample size (N). Totals: IHC $N = 159$, OHC $N = 529$, overall $N = 688$.

| KO_WT | Cell | Pair (level1–level2) | n1/n2 | mean1 | mean2 | Δ (95% CI) | $g$ | $p_{adj}$ (Holm) | Status |
|---|---|---|---|---|---|---|---|---|---|
| WT | IHC | Base–Middle | 26/28 | 0.702 | 0.241 | $+0.461\,(0.342, 0.580)$ | 2.16 | $6.27\times10^{-9}$ | Significant |
| WT | IHC | Middle–Apex | 28/27 | 0.241 | 0.173 | $+0.068\,(0.002, 0.133)$ | 0.55 | 0.0429 | Significant |
| WT | IHC | Base–Apex | 26/27 | 0.702 | 0.173 | $+0.528\,(0.411, 0.646)$ | 2.51 | $3.98\times10^{-10}$ | Significant |
| WT | OHC | Base–Middle | 81/85 | 0.419 | 0.161 | $+0.259\,(0.227, 0.290)$ | 2.53 | $2.57\times10^{-34}$ | Significant |
| WT | OHC | Middle–Apex | 85/100 | 0.161 | 0.123 | $+0.038\,(0.013, 0.063)$ | 0.45 | 0.00258 | Significant |
| WT | OHC | Base–Apex | 81/100 | 0.419 | 0.123 | $+0.297\,(0.266, 0.327)$ | 2.99 | $6.39\times10^{-41}$ | Significant |
| KO | IHC | Base–Middle | 26/26 | 0.512 | 0.329 | $+0.183\,(0.116, 0.250)$ | 1.51 | $6.00\times10^{-6}$ | Significant |
| KO | IHC | Middle–Apex | 26/26 | 0.329 | 0.287 | $+0.042\,(-0.062, 0.146)$ | 0.23 | 0.414 | Not significant |
| KO | IHC | Base–Apex | 26/26 | 0.512 | 0.287 | $+0.225\,(0.113, 0.337)$ | 1.11 | $4.34\times10^{-4}$ | Significant |
| KO | OHC | Base–Middle | 77/88 | 0.277 | 0.258 | $+0.0186\,(-0.0138, 0.0511)$ | 0.18 | 0.258 | Not significant |
| KO | OHC | Middle–Apex | 88/98 | 0.258 | 0.140 | $+0.118\,(0.088, 0.148)$ | 1.12 | $1.32\times10^{-12}$ | Significant |
| KO | OHC | Base–Apex | 77/98 | 0.277 | 0.140 | $+0.137\,(0.103, 0.171)$ | 1.20 | $1.07\times10^{-12}$ | Significant |

**Table 10.** Pairwise tonotopic contrasts for *normalized intensity* by genotype (WT/KO) and cell type (IHC/OHC). We report sample sizes ($n_1/n_2$), group means (mean1/mean2), difference $\Delta = \text{mean}_1 - \text{mean}_2$ with 95% CI, Hedges' $g$, and Holm-adjusted $p$-values. Positive Δ indicates level1 > level2.

| Block 1 | | | Block 2 | | | Block 3 | | |
|---|---|---|---|---|---|---|---|---|
| # | Manual | VASCilia | # | Manual | VASCilia | # | Manual | VASCilia |
| 1 | 112.88° | 111.10° | 12 | 99.06° | 95.34° | 23 | 93.13° | 91.16° |
| 2 | 109.81° | 110.56° | 13 | 92.32° | 95.23° | 24 | 94.97° | 94.14° |
| 3 | 78.61° | 76.35° | 14 | 104.65° | 102.91° | 25 | 81.81° | 81.49° |
| 4 | 101.31° | 95.34° | 15 | 92.07° | 92.69° | 26 | 83.60° | 87.44° |
| 5 | 94.44° | 97.28° | 16 | 91.53° | 87.61° | 27 | 91.46° | 90.36° |
| 6 | 93.99° | 93.89° | 17 | 99.88° | 99.33° | 28 | 91.01° | 90.81° |
| 7 | 98.62° | 94.76° | 18 | 77.42° | 75.48° | 29 | 89.46° | 89.60° |
| 8 | 87.31° | 88.01° | 19 | 100.13° | 97.38° | 30 | 92.29° | 92.47° |
| 9 | 88.23° | 90.00° | 20 | 91.15° | 89.55° | 31 | 104.42° | 102.40° |
| 10 | 99.05° | 98.39° | 21 | 92.70° | 91.60° | 32 | 90.53° | 88.45° |
| 11 | 113.96° | 113.20° | 22 | 83.06° | 86.16° | 33 | 88.57° | 90.35° |
| **Summary —** Mean (Manual/VASCilia): 94.04 / 93.35; Median: 92.31 / 92.47; SD: 8.83 / 8.44 | | | | | | | | |

**Table 11.** Per-cell orientation angles measured manually in Fiji and compared with VASCilia (related to Fig. 13).

| Block1 | | | Block2 | | |
|---|---|---|---|---|---|
| # | Manual | VASCilia | # | Manual | VASCilia |
| 1 | 103.75° | 99.68° | 10 | 91.18° | 90.53° |
| 2 | 87.73° | 89.43° | 11 | 85.14° | 84.44° |
| 3 | 88.17° | 87.95° | 12 | 89.42° | 86.52° |
| 4 | 72.48° | 80.68° | 13 | 80.01° | 85.75° |
| 5 | 103.04° | 98.58° | 14 | 94.86° | 89.51° |
| 6 | 79.96° | 76.09° | 15 | 89.39° | 88.89° |
| 7 | 84.80° | 85.24° | 16 | 82.46° | 84.13° |
| 8 | 85.56° | 80.89° | 17 | 91.14° | 85.75° |
| 9 | 85.87° | 82.95° | | | |
| **Summary —** Mean (Manual/VASCilia): 87.94 / 86.88; Median: 87.73 / 85.75; SD: 7.80 / 5.90 | | | | | |

**Table 12.** Comparison of orientation angles measured manually in Fiji and by VASCilia, displayed in two horizontal panels (related to Fig. 14).

| Block A | | | Block B | | | Block C | | |
|---|---|---|---|---|---|---|---|---|
| # | Manual | VASCilia | # | Manual | VASCilia | # | Manual | VASCilia |
| 1 | 1.603 | 1.696 | 6 | 2.252 | 2.179 | 11 | 1.989 | 1.986 |
| 2 | 1.629 | 1.715 | 7 | 2.086 | 2.009 | 12 | 1.703 | 1.741 |
| 3 | 1.855 | 1.858 | 8 | 2.832 | 2.816 | 13 | 1.313 | 1.277 |
| 4 | 1.711 | 1.634 | 9 | 2.035 | 1.965 | 14 | 1.635 | 1.787 |
| 5 | 1.193 | 1.218 | 10 | 2.216 | 2.249 | 15 | 1.333 | 1.400 |
| | | | | | **Mean / SD** | **Mean** | **1.826** | **1.835** |
| | | | | | | **Std Dev** | **0.426** | **0.405** |
| | | | | | **Paired $t$-test $p$-value** | | **0.609** | |
| | | | | | **Wilcoxon signed-rank $p$-value** | | **0.720** | |

**Table 13.** Per-cell length values from human-annotated ground truth for *Eps8* KO mice using Fiji and VASCilia for 15 cells. The comparison showed no statistically significant difference between the two methods. Specifically, the mean stereocilia length was 1.826 µm for Fiji and 1.835 µm for VASCilia, with standard deviations of 0.426 µm and 0.405 µm, respectively. A Wilcoxon signed-rank test and a paired t-test both confirmed the absence of significant differences.

| Block A | | | Block B | | | Block C | | |
|---|---|---|---|---|---|---|---|---|
| # | Manual | VASCilia | # | Manual | VASCilia | # | Manual | VASCilia |
| 1 | 1.964 | 1.948 | 6 | 1.777 | 1.803 | 11 | 1.658 | 1.684 |
| 2 | 1.873 | 1.879 | 7 | 1.756 | 1.790 | 12 | 1.430 | 1.248 |
| 3 | 2.139 | 2.105 | 8 | 2.754 | 2.739 | 13 | 1.186 | 1.082 |
| 4 | 2.260 | 2.356 | 9 | 1.475 | 1.592 | 14 | 0.866 | 0.886 |
| 5 | 1.735 | 1.955 | 10 | 2.117 | 1.948 | 15 | 1.697 | 1.751 |
| | | | | | **Mean / SD** | **Mean** | **1.779** | **1.784** |
| | | | | | | **Std Dev** | **0.455** | **0.468** |
| | | | | | **Paired $t$-test $p$-value** | | **0.851** | |
| | | | | | **Wilcoxon signed-rank $p$-value** | | **0.639** | |

**Table 14.** Per-cell length values from human-annotated ground truth for $Cdh23^{-/-}$ mice using Fiji and VASCilia for 15 cells. The comparison showed no statistically significant difference between the two methods. Specifically, the mean stereocilia length was 1.779 µm for Fiji and 1.784 µm for VASCilia, with standard deviations of 0.455 µm and 0.468 µm, respectively. A Wilcoxon signed-rank test and a paired t-test both confirmed the absence of significant differences.

| Block A | | | Block B | | | Block C | | |
|---|---|---|---|---|---|---|---|---|
| **ID** | **Manual** | **VASCilia** | **ID** | **Manual** | **VASCilia** | **ID** | **Manual** | **VASCilia** |
| cell8 | 86.32° | 88.17° | cell13 | 88.32° | 90.00° | cell18 | 101.30° | 101.31° |
| cell7 | 83.26° | 86.68° | cell14 | 102.46° | 96.71° | cell20 | 104.85° | 109.03° |
| cell24 | 95.98° | 92.49° | cell15 | 60.09° | 68.20° | cell48 | 140.98° | 139.09° |
| cell6 | 90.68° | 90.00° | cell9 | 109.07° | 103.39° | cell33 | 140.20° | 138.99° |
| cell5 | 91.56° | 91.61° | cell11 | 119.81° | 117.98° | cell26 | 107.27° | 104.04° |
| cell4 | 87.22° | 89.26° | cell44 | 113.11° | 112.99° | cell29 | 122.89° | 123.69° |
| cell1 | 71.19° | 75.23° | cell45 | 135.39° | 131.28° | cell32 | 77.60° | 77.91° |
| cell3 | 85.28° | 84.35° | cell42 | 92.95° | 92.39° | cell38 | 29.47° | 31.18° |
| cell2 | 74.90° | 79.00° | cell34 | 98.97° | 96.77° | cell36 | 80.41° | 82.03° |
| cell39 | 129.93° | 125.54° | cell35 | 107.80° | 103.24° | cell40 | 126.73° | 128.88° |
| cell27 | 146.07° | 151.19° | cell30 | 121.94° | 121.61° | cell37 | 117.13° | 115.46° |
| cell31 | 123.36° | 128.99° | cell19 | 119.58° | 124.22° | cell47 | 64.80° | 66.80° |
| cell28 | 90.53° | 90.00° | cell16 | 90.43° | 88.98° | cell41 | 116.55° | 114.90° |
| cell25 | 110.79° | 109.44° | cell21 | 92.18° | 92.39° | cell43 | 122.59° | 122.35° |
| cell22 | 151.15° | 149.66° | cell12 | 75.11° | 78.69° | cell46 | 107.57° | 106.86° |
| cell17 | 136.45° | 135.00° | | | | | | |
| | | | | | | **Mean** | **103.090°** | **103.222°** |
| | | | | | | **Std Dev** | **24.940** | **24.061** |
| | | | | | | **Paired $t$-test $p$-value** | **0.783** | |
| | | | | | | **Wilcoxon signed-rank $p$-value** | **0.965** | |

**Table 15.** Per-bundle orientation values (degrees) for a PCP-deficit mouse cochlear dataset[41], see Fig. 23, measured in Fiji (manually) and with VASCilia (automated). A paired $t$-test and a Wilcoxon signed-rank test indicate no significant difference.

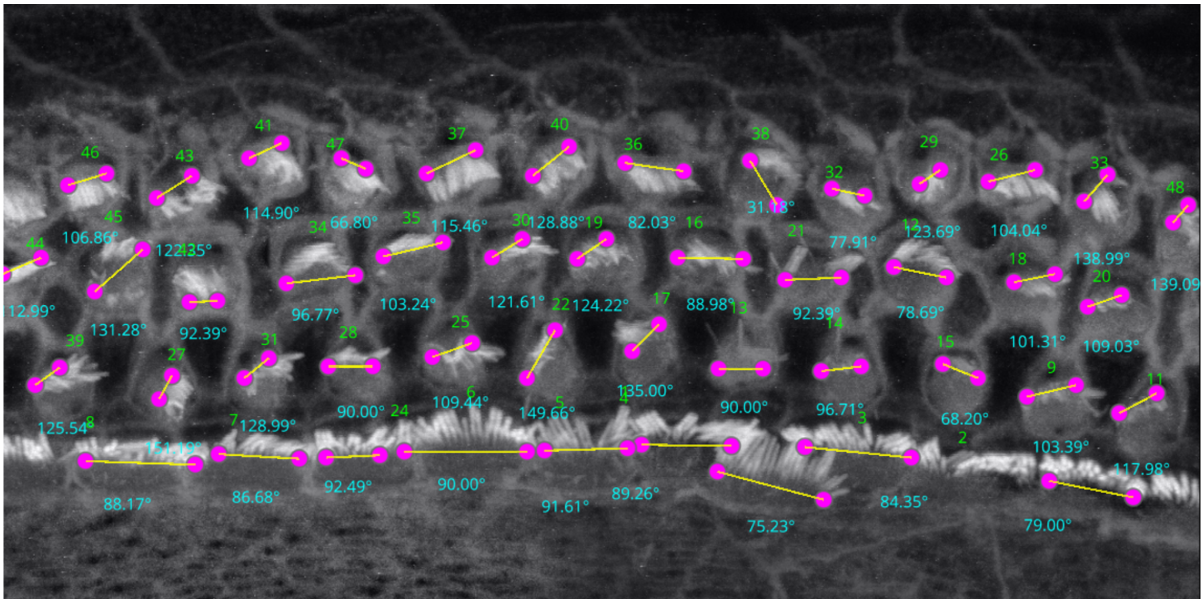


**Figure 23.** Angle computation on a PCP-deficit cochlear dataset[41]. VASCilia recovers bundle orientation and agrees closely with the manual measurements using Fiji (see Table 15; paired $t$-test $p = 0.783$, Wilcoxon $p = 0.965$).

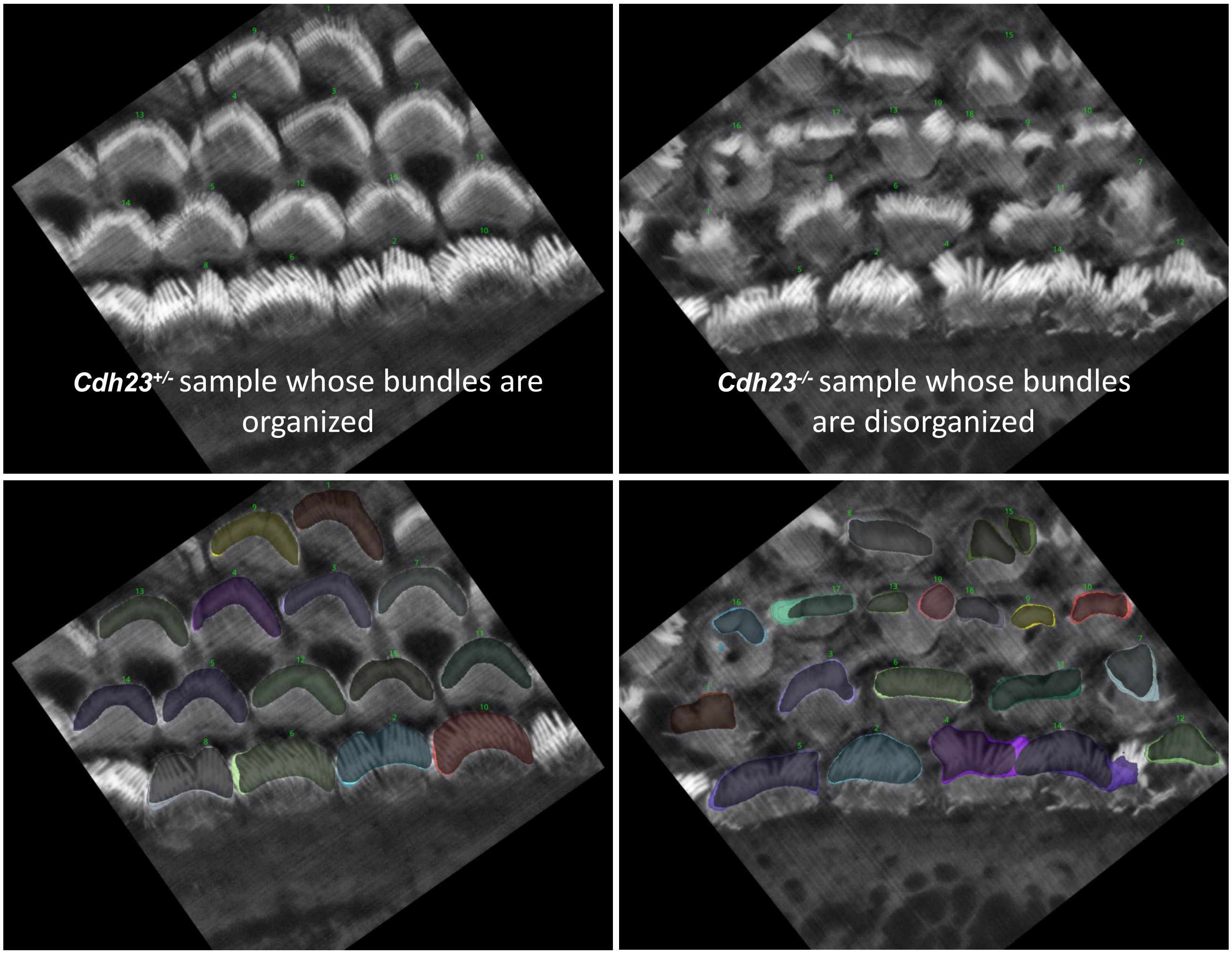


**Figure 24.** Representative images for *Cdh23*$^{+/-}$ and *Cdh23*$^{-/-}$ mouse

**VASCilia analysis for Lab A's datasets: screenshots for various samples.**

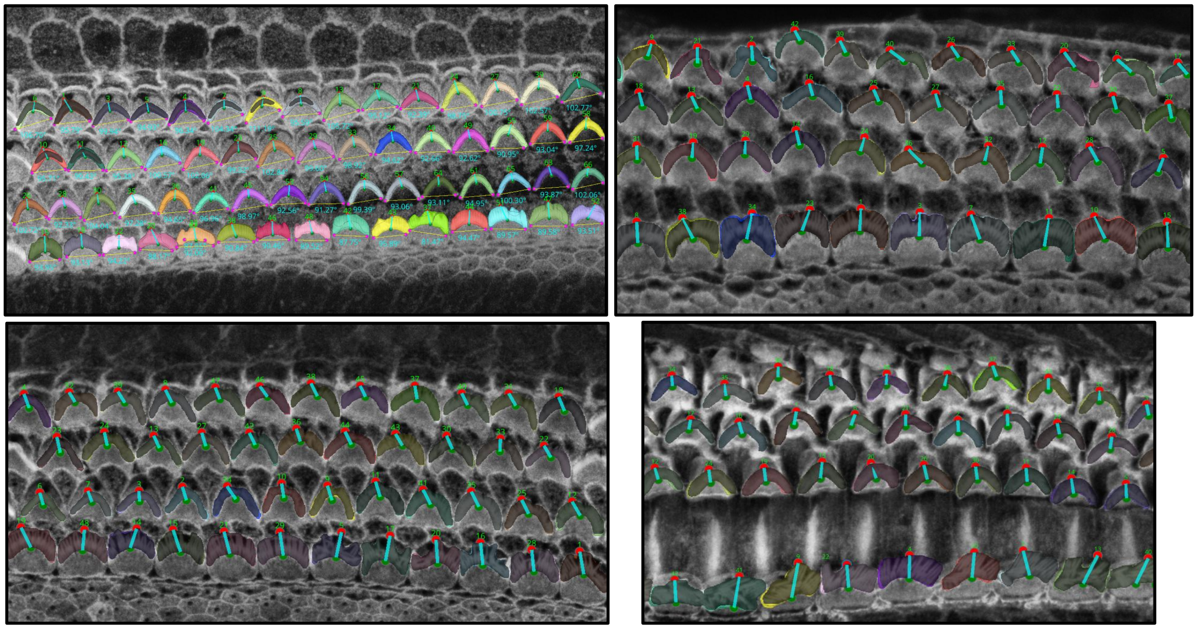

**Figure 25.** Screenshot of the software with different samples.

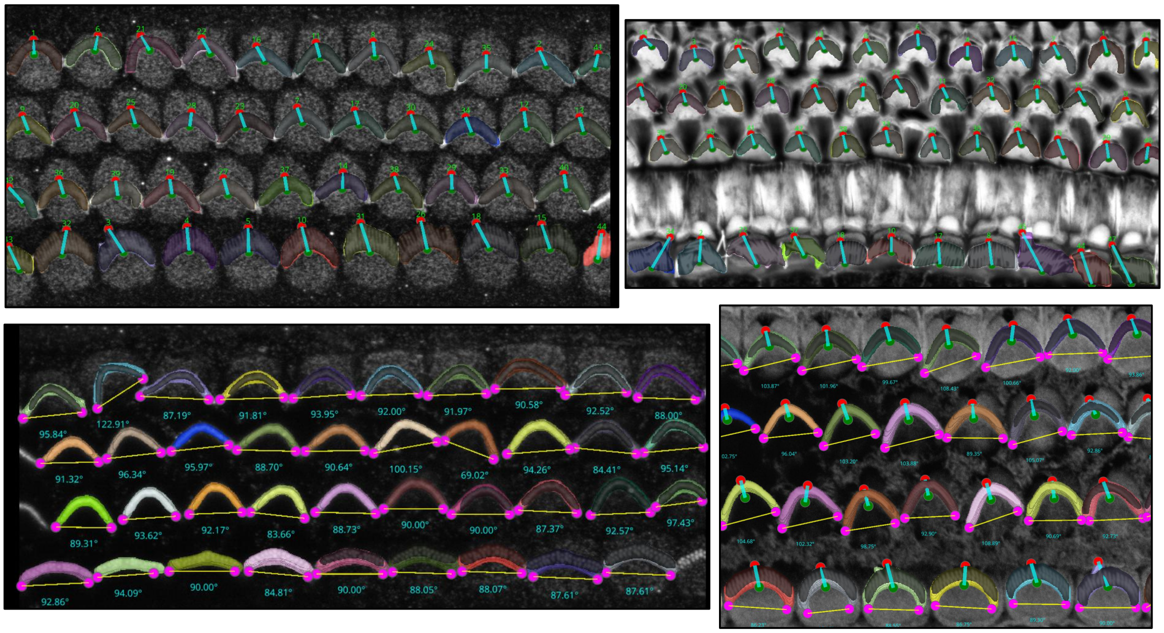

**Figure 26.** Screenshot of the software with different samples.

### VASCilia analysis for Lab B's datasets: screenshots for three samples.

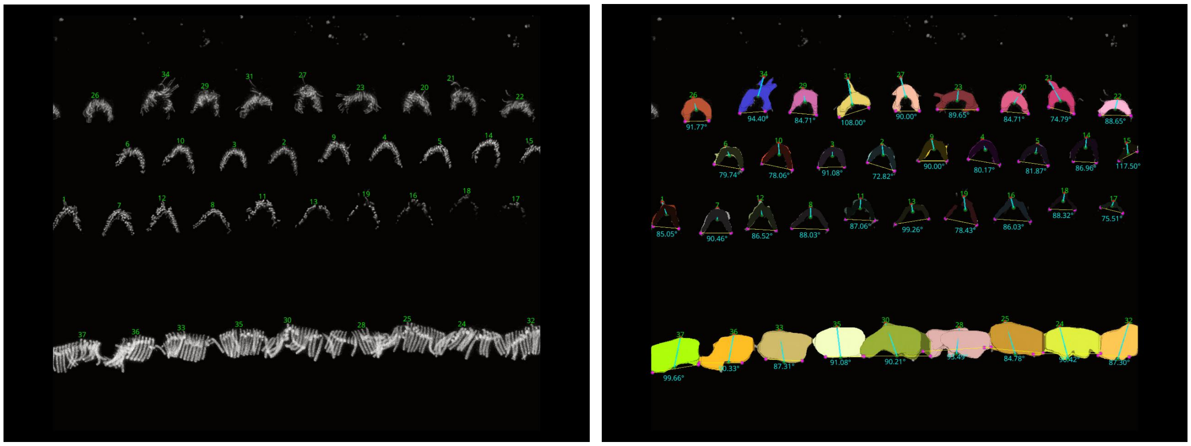

**Figure 27.** Screenshot of the software with Lab B's mouse cochlea data.

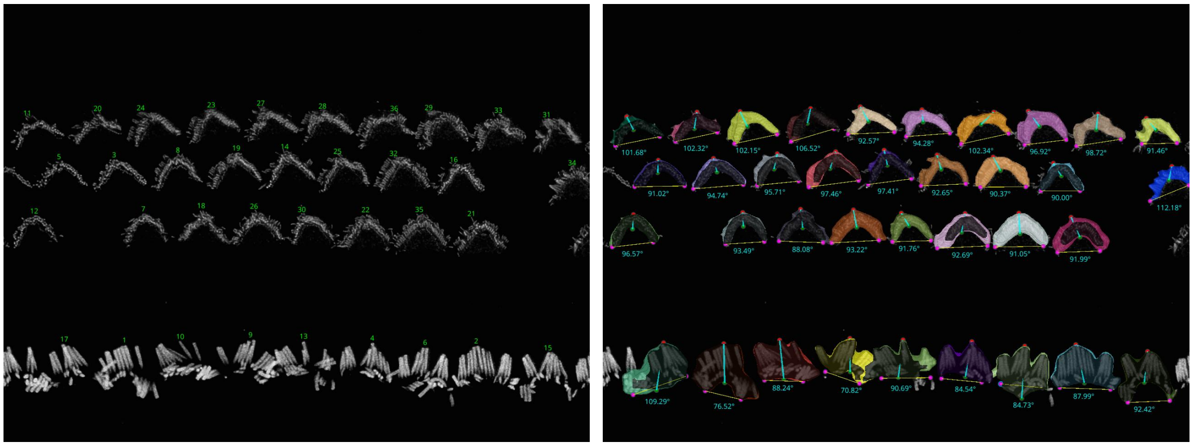

**Figure 28.** Screenshot of the software with Lab B's mouse cochlea data.

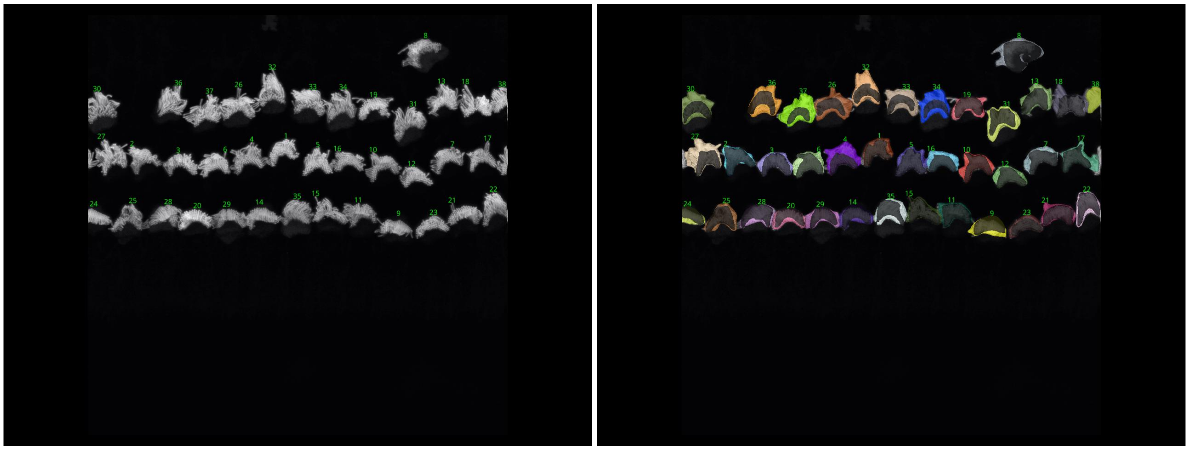

**Figure 29.** Screenshot of the software with Lab B's human cochlea data.

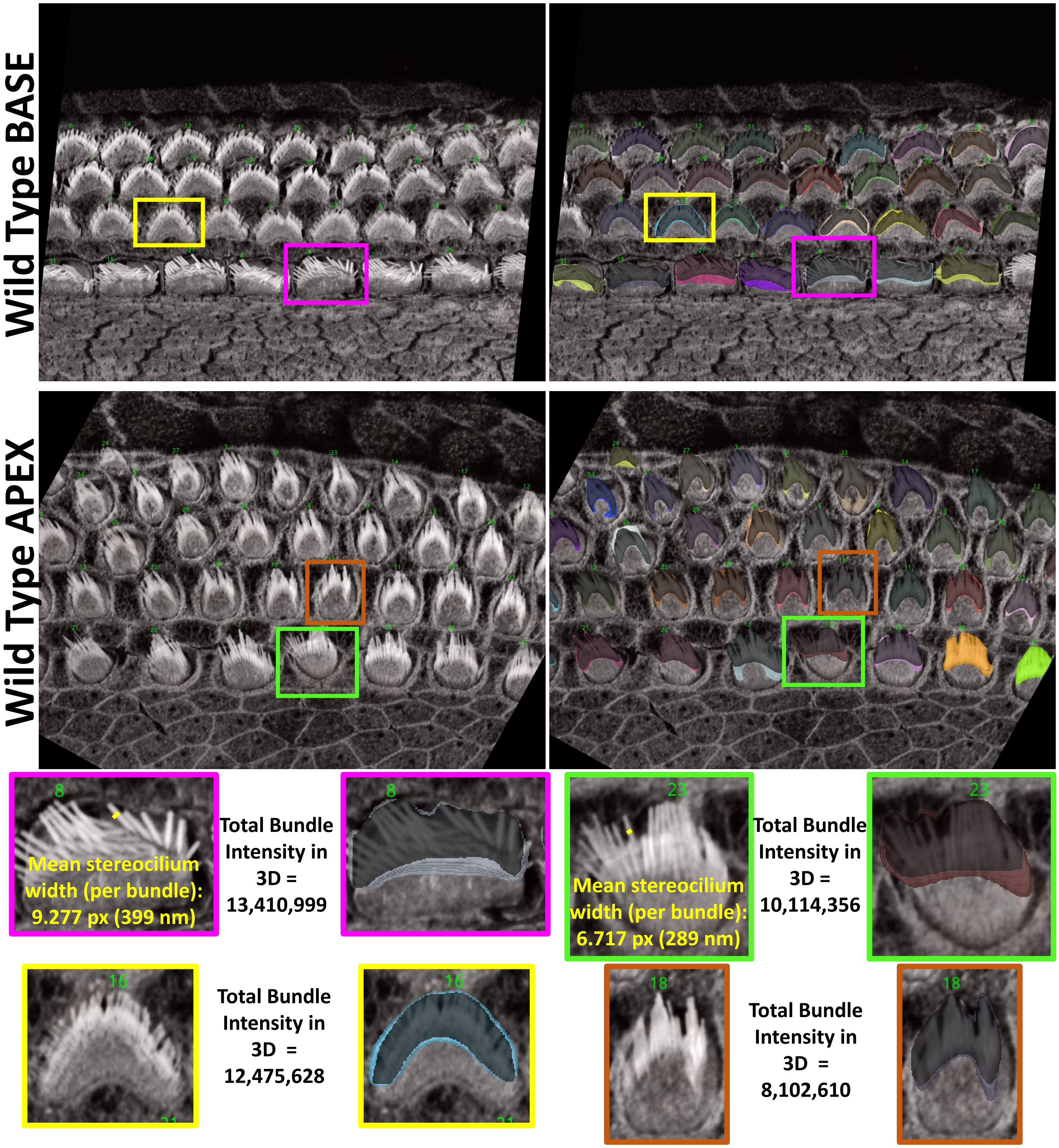


**Figure 30.** Visual comparison of base and apex bundles in WT animals reveals that the base appears stiffer, with wider stereocilia and bundles, while the apex displays finer stereocilia and tighter bundles, leading to a reduced phalloidin intensity signal.
